# Flexible cognition in context-modulated reservoir networks

**DOI:** 10.1101/2022.05.09.491102

**Authors:** Nicolas Y. Masse, Matthew C. Rosen, Doris Y. Tsao, David J. Freedman

**Author notes:** Equal contribution (Nicolas Y. Masse, Matthew C. Rosen).

## Abstract

The brains of all animals are plastic, allowing us to form new memories, adapt to new environments, and to learn new tasks. What is less clear is how much plasticity is required to perform these cognitive functions: does learning require widespread plasticity across the brain, or can learning occur with more rigid networks, in which plasticity is highly localized? Here, we use biologically-inspired recurrent neural network (RNN) models to show that rapid multitask learning can be accomplished in reservoir-style networks, in which synaptic plasticity is sparse and highly localized. Crucially, only RNNs initialized with highly specific combinations of network properties, such as topology, normalization and reciprocal connection strength, are capable of such learning. Finally, we show that this rapid learning with localized plasticity can be accomplished with purely local error signals, without backpropagation, using a reinforcement learning setup. This work suggests that rapid learning in artificial (and potentially biological) agents can be accomplished with mostly-rigid networks, in which synaptic plasticity is highly constrained.

## Introduction

An organism’s ability to learn – that is, to adjust its behavior in response to events it experiences and stimuli it encounters in its environment – is an extraordinarily powerful adaptive advantage: the more flexibly an organism can learn, the better equipped it is to survive changes, either to the environment or to itself. Understanding the cellular basis of this ability is one of the central goals of systems neuroscience.

A primary candidate mechanism for learning in the brain is the plasticity of synaptic connections between neurons, involving changes either to their strength or their existence. Beginning in development, synaptic plasticity plays a critical role in regulating how information flows through circuits of neurons, which governs the set of computations those circuits can perform [1]. This developmental plasticity is also generally necessary for survival: genetic or biochemical perturbations that arrest or diminish synaptic plasticity during development are associated with a myriad set of diseases and dysfunctions [2, 3]. Even after this early critical period, however, the adult brain still maintains an impressive capacity for plasticity, experimentally confirmed by studies employing behavioral assays, pharmacological and genetic inactivation, anatomical lesions, and *in vivo* and *in vitro* electrophysiology [4–6].

However, the manner in which learning engages this substantial capacity for plasticity during adulthood remains unclear. Monitoring synapses *in vivo* is technically challenging, and it is currently not feasible to directly measure synaptic connections throughout the brain across the longer time scales of many forms of learning [7, 8]. Only recently have technically innovative studies in small animals documented how specific synaptic connections are altered during task learning [9–11]. But despite these new (yet sparse) experimental results, it is still not clear whether task learning requires widespread synaptic changes throughout the brain, or whether learning can be accomplished with more constrained and localized plasticity. That said, there are several reasons to believe that limiting task-related synaptic plasticity during learning might be advantageous. First, long term synaptic potentiation (LTP) is metabolically expensive [12]. Second, methods to constrain synaptic plasticity when learning tasks sequentially can reduce interference between synaptic weights learned for different tasks, mitigating catastrophic forgetting of previously-acquired knowledge [13–15]. Third, the most effective known algorithm to alter synaptic connections throughout the network during task learning is backpropagation, and despite many theoretical advances [16–18], backpropagation throughout the brain and through time would be challenging for biological circuits to implement. Thus, while the adult brain is capable of widespread plasticity, it is possible that more constrained plasticity underlies many forms of learning.

Here, we use biologically-inspired RNN models to ask how plastic a neural network needs to be to support the learning of cognitive tasks [19, 20]. These models, trained using backpropagation through time, can learn an impressive array of complex tasks, and offer the opportunity for detailed examination of network activity and circuit structure which support task performance [21–23]. Drawing inspiration from earlier generations of model, such as echo state networks and [24] liquid state machines [25], as well as later variants like FORCE training [26], we show that rather than relying on widespread changes in synaptic weights throughout the network during training, RNNs in which synaptic plasticity is constrained to be sparse and highly localized can rapidly and robustly learn multiple tasks. Specifically, we only train connections in the output layer and in a top-down layer that linearly transforms the task context. This top-down layer projects onto the recurrent layer, altering its neural dynamics and allowing it to flexibly perform multiple tasks involving working memory, decision making, and categorization. Crucially, only specific combinations of hyperparameters that determine network properties (topology, normalization, etc.) produce RNNs capable of rapid and robust learning. Finally, we demonstrate that learning can be performed with purely local reward signals using a reinforcement learning setup between two cooperating actors, eliminating the need to backpropagate error signals through the network and through time, which may be challenging to implement in biological circuits [17, 27, 28]. This work suggests that sparse and localized synaptic changes can support rapid and biologically-plausible learning of multiple tasks, pointing to novel machine learning approaches and architectures for rapid and robust learning and motivating novel hypotheses to be tested by neuroscience experiments.

## Results

The goal of this study was to determine whether rapid multitask learning in recurrent network models can be accomplished with minimal and highly localized synaptic plasticity. We first demonstrate how this can be performed using supervised learning, and then go on to show how this can be accomplished in a more biologically-plausible manner using reinforcement learning and only local reward signals.

We trained RNNs to learn six tasks (Figure 1A) used in cognitive and systems neuroscience experiments, which require perceptual and categorical decisions, as well as short-term memory and cognitive control [29, 30] (see Methods). The stimulus in all six tasks was visual motion direction, presented in one of six directions. In all tasks, the network was required to maintain “fixation” before selecting the correct action during the test period. All tasks were learned simultaneously, and the identity of the task was cued to the network. In order to select the correct action during any task, networks needed to maintain and manipulate information in working memory, and compare stimuli presented at different times based on their motion direction or category membership. Importantly, while the stimuli were similar across tasks, the correct response differed, forcing the network to utilize the task identity to accurately perform the task.

**Figure 1:**
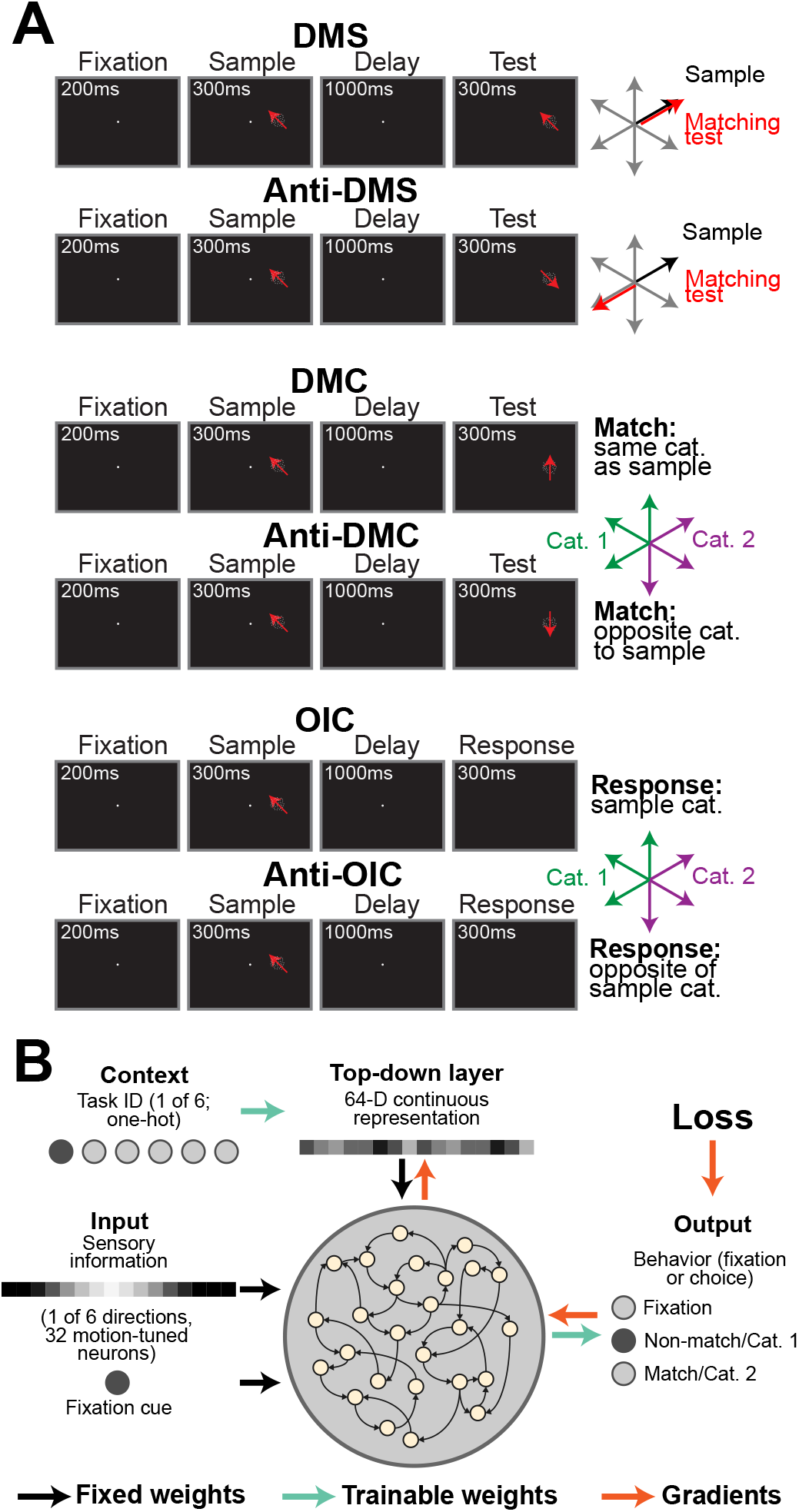
Cognitive tasks and network architecture. **A)** Networks were trained on six different cognitive tasks. In the delayed match-to-sample (DMS) and anti-DMS tasks (rows 1 and 2), the network was rewarded for indicating whether the test stimulus direction matched (was rotated 180° away from) the sample stimulus direction. In the delayed match-to-category (DMC) and anti-DMC tasks (rows 3 and 4), the network was rewarded for indicating whether the sample direction belonged to the same (opposite) category as the test direction. The two categories are indicated by green and purple arrows. In the one interval category (OIC) and anti-OIC tasks (rows 5 and 6), the network was rewarded for indicating which category the sample stimulus belonged to. The correct response for the OIC task was opposite that for the anti-OIC task. **B)** The network consisted of several layers whose weights were either fixed (black arrows) or trainable (green arrows). Training the top-down context layer required backpropagating the gradient from the output, and through the recurrent layer (red arrows).

Our biologically-inspired RNNs, similar to those in previous studies [21–23], consisted of several layers (Figure 1B). Since we were interested in whether networks can rapidly learn multiple tasks with localized synaptic changes, we limited weight changes to the output layers and the top-down layer which linearly transformed the task identity (green arrows, Figure 1B). Weights in all other layers were kept fixed and were not adjusted during training (black arrows). The trainable weights consisted of ∼ 0.1% of all network parameters. Training still required backpropagation through time and through hidden layers, as the gradient of the loss with respect to the top-down layer needed to propagate through the recurrent layer (red arrows). We demonstrate that networks can learn the tasks without the need for backpropagation later on (Figure 4). Full network architecture and training procedure is detailed in the Methods.

### Searching the space of network properties

Given that synaptic plasticity during development is crucial for establishing adult brains with properly functioning neural circuits needed to support behavior and learning [1–3], we hypothesized that only RNNs with specific characteristics would be capable of rapidly learning all six tasks with limited synaptic plasticity. To test this, we defined a set of 26 hyperparameters, based on physical properties defining neural circuits *in vivo*, that governed the connectivity within our model. These hyperparameters controlled the strength of connections between and within excitatory and inhibitory neurons in the recurrent layer, their connections with the bottom-up and top-down layers, network topology, and normalization, amongst other attributes (see Methods).

Since we did not know *a priori* which hyperparameters would produce RNNs capable of rapid learning, we performed a random search across these 26 hyperparameters, where we would 1) randomly sample the 26 hyperparameters from defined ranges, 2) randomly sample network weights from probability distributions governed by these 26 hyperparameters, 3) run the resulting RNN on a batch of trials, and 4) if the mean RNN activity was not excessively strong, rapidly train the output and top-down layers on 150 batches of 256 trials (6,400 trials per task). In comparison, Yang *et al*. [21] trained RNNs for 150,000 trials per task.

Out of the 10,114 RNNs we randomly sampled using this procedure, only a fraction (2,725, or 26.9%) generated mean activity below 1 (shaded region, Figure 2A), which we empirically observed was strongly associated with successful training (networks with high activity became unstable). RNNs with mean activity above this value were no longer considered. Since all tasks required maintaining information in short-term memory, we then calculated how accurately the RNN could encode the identity of the sample stimulus at the end of the delay period (see Methods, Figure 2B). The distribution of the sample decoding accuracies, measured at the end of the delay period, was centered near chance level (1/6), indicating that most RNNs were not capable of maintaining sample information across the delay period. This is unsurprising, as the networks were not trained to encode information in short-term memory. However, the sample decoding accuracy from a small subset of RNNs was closer to one (perfect decoding), suggesting that some RNNs were innately capable of reliably maintaining information needed to solve the tasks in the form of persistent activity during the delay period, without any training to do so.

**Figure 2:**
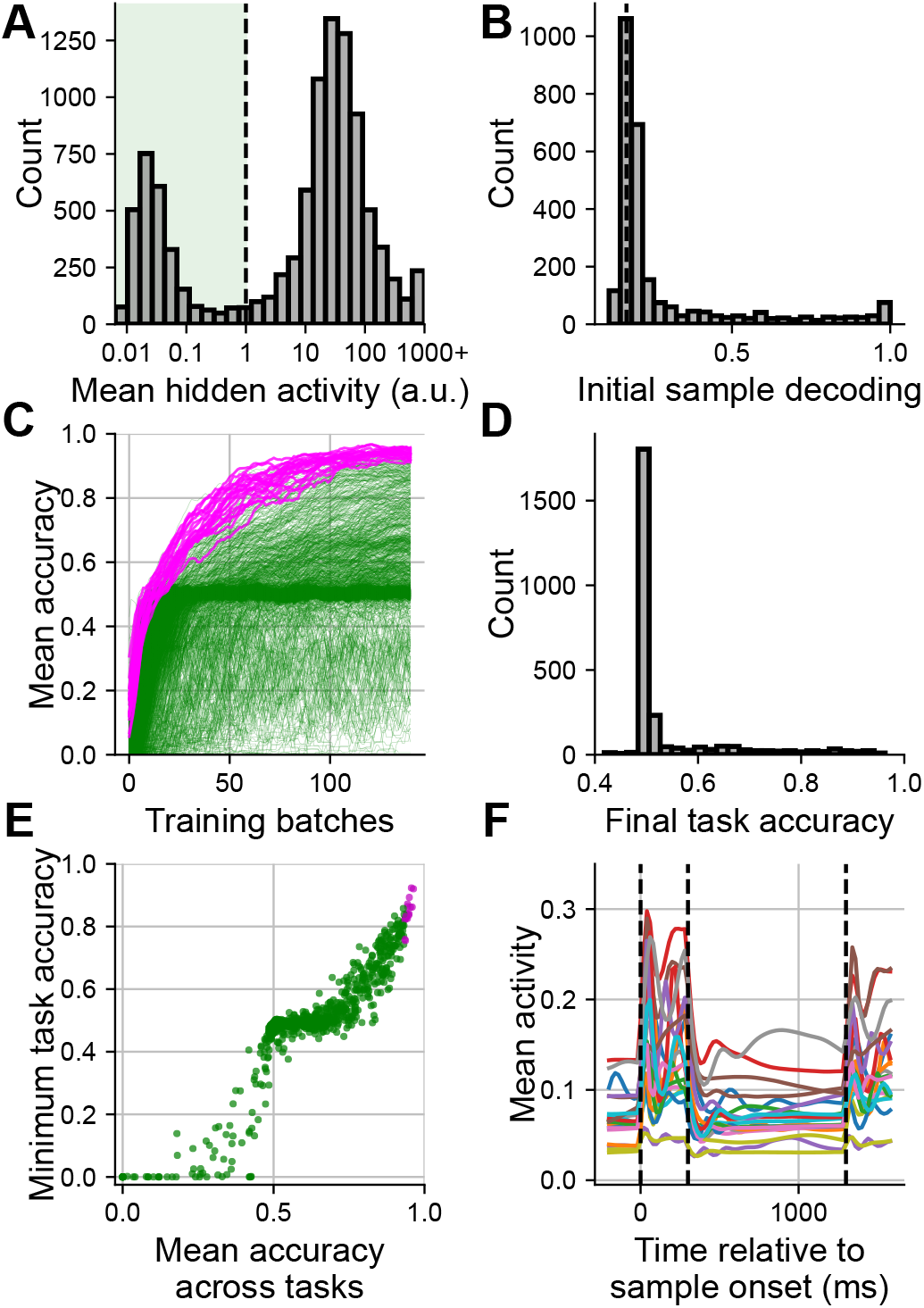
Properties of the 10,114 RNNs trained on the six cognitive tasks. **A)** Mean activity from the recurrent layer measured before training. Networks with mean activity above 1 (indicated by the vertical dashed line) were discarded. **B)** Sample decoding accuracy measured at the end of the delay period and before network training. Chance accuracy (1/6) is indicated by the vertical dashed line. **C)** Mean accuracy across the six tasks measured after each training batch for the 2,725 RNNs with suitable mean activity (green and magenta traces). The traces were smoothed, with the accuracy averaged across the last 10 batches. The magenta traces are from the 20 highest-performing RNNs. **D)** Distribution of the mean accuracy across all six tasks at the end of training. The accuracy of 59/2,725 RNNs was <40%, and are not shown. **E)** Scatter plots showing the mean (x-axis) and minimum (y-axis) accuracies across the 6 tasks. Magenta circles indicate the 20 high-performing RNNs. **F)** Mean activity from the recurrent layer across trial time for the 20 high-performing RNNs. Sample onset/offset, and test onset are indicated by the dashed vertical lines.

We then trained each of the 2,725 RNNs with suitable mean activity on 150 batches of 256 trials. The mean accuracy across the six tasks during training (Figure 2C) and the distribution of accuracies at the end of training (Figure 2D) show that most RNNs achieved accuracies of ∼ 50%. This is the level of performance expected if the RNNs learn to correctly maintain fixation, but not how to correctly respond during the test epoch. However, a small fraction of RNNs were capable of rapidly learning the six tasks simultaneously to a reasonably high accuracy: 56/2,725 RNNs achieved >90%, and 4/2,725 achieved >95%. The traces from the top 20 RNNs, shown in magenta (Figure 2C), all achieved >93%. To ensure that these RNNs’ high performance reflected learning of all tasks, we compared the mean accuracy across all six tasks (x-axis, Figure 2E) to the minimum accuracy across the six tasks (y-axis). For all but one of the 20 high-performing RNNs (magenta circles), the minimum observed task accuracy exceeded 80%, suggesting that these RNNs learned to competently perform all tasks. We also examined the mean network activity across time (measured from the recurrent layer during the DMS task) of these high-performing RNNs (Figure 2F). We observed a large variety of neural responses, highlighting how different sets of hyperparameters generate distinct patterns of neural activity. Activity traces from many of the RNNs resembled what is typically found *in vivo*, with transient and sustained responses to the sample and test stimuli, and some ramping activity during the delay period, suggesting that neural dynamics of these networks, at some level, bear some similarity to neural dynamics found *in vivo*.

### Reliability and generalization of high-performing network hyperparameters

These results show that RNNs are capable of rapidly learning multiple tasks with highly localized synaptic changes, provided that they are properly initialized. However, given that we trained thousands of RNNs, these high-performing RNNs may be the result of sample bias, rather than robust initialization for multitask learning. To test whether the hyperparameters that defined the 20 highest-performing RNNs in our initial sweep could reliably generate high-performing networks, we repeatedly resampled and retrained network weights from these hyperparameters, generating 5 new RNNs for each set. We then compared task accuracy of the 5 resampled and retrained RNNs (y-axis, Figure 3A) with the original accuracies obtained in the large-scale sweep from Figure 2 (x-axis). For 11/20 of the high-performing hyperparameter sets (the 20 high-performing sets are indicated by the magenta circles), task accuracy of the 5 resampled and retrained RNNs was consistent (within 5 percentage points on average) with the original accuracies obtained in the large-scale sweep. Going forward, we refer to these 11 hyperparameter sets as “reliable” (see Figure 4). For another 6 hyperparameter sets, mean accuracy after resampling and retraining decreased, but was still >80%, and for the remaining 3 hyperparameter sets, task accuracy of the resampled RNNs was lower. Thus, while some hyperparameter sets achieved high accuracy due to selection bias during the initial sweep, most hyperparameters that generated high-performing RNNs in the initial sweep do so consistently, even after randomly resampling their connection weights.

**Figure 3:**
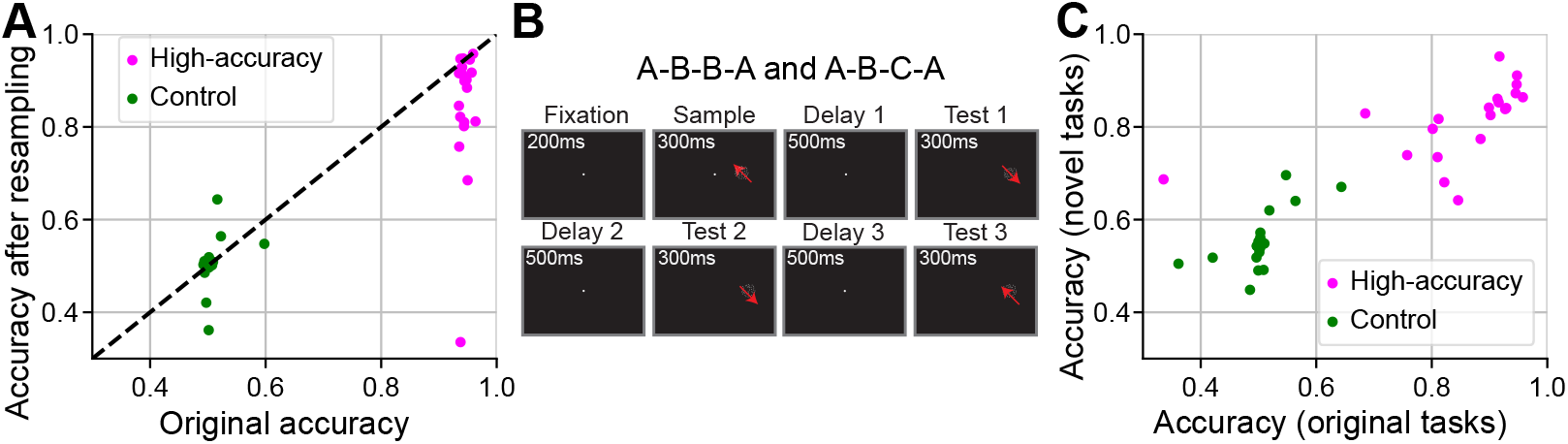
Reliability and generalization of high-performing network hyperparameters. **A)** Scatter plot showing mean accuracy across the six tasks from the original sweep (x-axis) against the mean accuracy after resampling network hyperparameters and retraining (y-axis). Magenta circles are from the 20 high-performing hyperparameter sets. The green circles are a randomly selected subset of hyperparameter sets that scored between 45 and 65% in the original sweep. **B)** We tested generalization using the A-B-B-A and A-B-C-A tasks. In both tasks, a sample stimulus was shown, followed by three test stimuli. The RNN had to determine whether the sample and test stimuli matched. In the A-B-B-A task, there was a 50% chance subsequent test stimuli had the same direction. For the A-B-C-A task, this probability was 0% **C)** Scatter plot using the same hyperparameter sets in A), showing the mean accuracy on the original six tasks after resampling and retraining (x-axis) against the mean accuracy on the two new tasks (y-axis).

**Figure 4:**
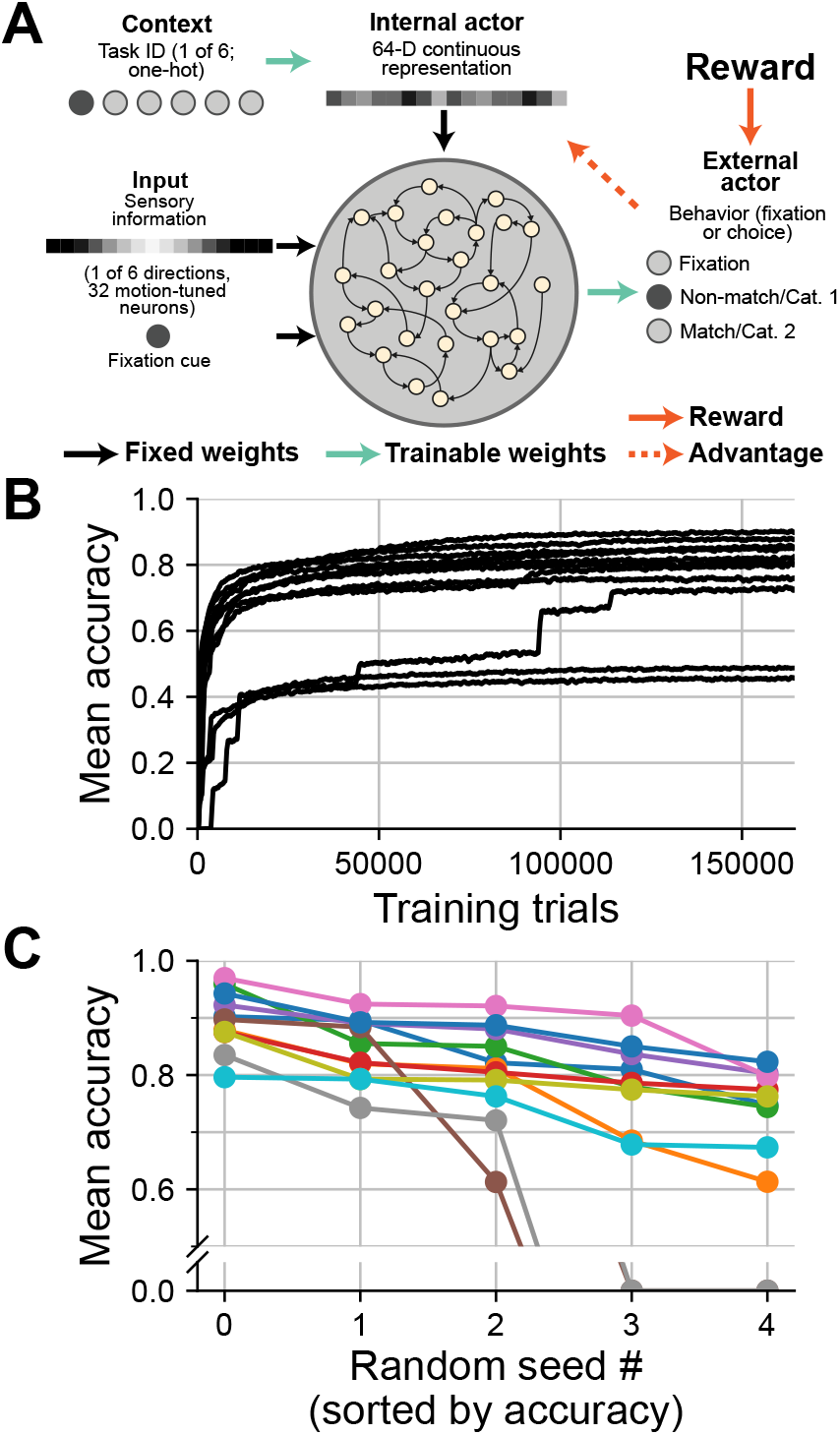
Training RNNs without backpropagation through time using RL. **A)** The network architecture was similar to the one shown in Figure 1B, but now consists of two actors trained using RL: the external actor acts on the external environment (e.g. responds to the tasks), and the internal actor acts on the internal environment (the recurrent layer). This removes the need for error signals to backpropagate through hidden layers (red arrows) or time. **B)** Mean trial accuracy across the six cognitive tasks for the 11 reliable high-performing hyperparameter sets during RL training. Each trace represents the average performance taken across 5 training runs per hyperparameter set, each with a unique random seed. **C)** Post-training accuracy of the 5 random seeds for each hyperparameter set. Seeds sorted in decreasing order of accuracy, and traces colored by hyperparameter set.

We repeated this analysis using an equal number of randomly selected network hyperparameters that achieved between 45 and 65% accuracy in our large-scale sweep (Figure 3A, green circles). None of these 20 hyperparameter sets achieved an accuracy >70% after resampling and retraining, further suggesting that successful rapid learning is limited to a small subset of network hyperparameters.

We also wanted to determine whether training the top-down weights was necessary to accurately learn all six tasks. Thus, we repeated our sweep used for Figure 2, except that top-down weights were randomized across tasks but frozen from initialization, and only output weights were trained (Figure S1). The top 20 hyperparameter sets from this sweep achieved lower mean accuracy across the six tasks (*p <* 10^−7^, Wilcoxon rank-sum test) and a lower minimum accuracy (*p <* 10^−7^), confirming that networks with trainable top-down weights are more capable of rapidly learning multiple tasks.

Next, we asked whether these high-performing networks could successfully generalize to novel tasks. Thus, we resampled network weights using the 26 hyperparameters of the same 20 high-performing networks, and the same 20 networks from the control group, and trained the resulting RNNs to perform two delayed match-to-sample tasks with multiple distractors: the A-B-B-A [31] and A-B-C-A [23] tasks (Figure 3B). In both tasks, a sample stimulus was followed by three sequentially presented test stimuli, and the RNN had to indicate whether each test matched the sample. In the A-B-B-A task, if a test stimulus was a non-match, there was a 50% probability that the test would be repeated immediately. This forced the RNN to encode sample and test stimuli in different ways: if the sample and test were encoded in the same manner, then the RNN would not be able to distinguish between a test that matched the sample compared with a repeated non-match [23]. In the A-B-C-A task, non-matching test stimuli were never repeated during a single trial, so the network was not required to represent sample and test stimuli in different formats. Importantly, the temporal sequence of events in A-B-B-A and A-B-C-A differs from that of the original six tasks, all of which shared a common temporal structure (in which the sample, delay, and test epochs were identically timed).

On average, the high-performing RNNs achieved 81.0% on the A-B-B-A task and 81.5% on the A-B-C-A task (magenta circles, Figure 3C), compared to 58.0% and 52.6% for the control group, respectively (black circles; *p <* 10^−7^, N = 20, Wilcoxon rank-sum test). Similar to the reliability test in Figure 3A, most, but not all, RNNs successfully learned the two tasks, with 13/20 achieving a combined accuracy of 80%, and 2 of those 13 exceeding 90%. Furthermore, a network’s ability to generalize to novel tasks was correlated with its ability to reliably solve the original task set (*R* = 0.79, *p <* 10^−4^, N = 20, Spearman’s rank correlation). Thus, many of the high-performing network hyperparameters generate RNNs that can reliably solve novel tasks.

### Biologically-plausible learning

So far, we have shown that RNNs can rapidly learn new tasks with synaptic changes localized to the output and top-down layers, but training still requires backpropagation through hidden layers and through time, which is hypothesized to be difficult for biological circuits to implement [17, 27, 28]. Thus, we wanted to determine whether RNNs could learn all six cognitive tasks in a more biologically-plausible manner, using only local error signals (i.e. without backpropagation). We made two key changes to this end (Figure 4A). First, we trained RNNs using policy-based reinforcement learning (RL) as opposed to supervised learning. In this setup, the output layer now contains a policy head, from which we sample which actions the networks perform, and a critic head, which estimates the discounted future reward (Figure 4A, see Methods). Second, we replaced the top-down layer with a second actor, also trained using RL. This second actor still receives as input the one-hot context signal, but its output now consists of a continuous-valued linear policy function, from which we sample actions that will “act” (i.e. linearly project) onto the recurrent layer. Thus, our setup now consists of two interacting RL-trained actors: one that observes and acts upon the external environment in the traditional manner, and a second that acts upon the “internal” environment (i.e. the recurrent layer). This, in principle, should allow the RNN to learn the proper top-down signal to control activity in the recurrent layer without the need for backpropagation of error signals.

We tested this method on new RNNs generated from the 11 high-performing hyperparameter sets identified as reliable in our followup to the large-scale sweep (Figure 3A). We trained these networks using RL for ∼ 160,000 trials per task. Given the variability in performance when training via RL [32], we trained RNNs from each hyperparameter set 5 times, using a different random seed each time. RNNs from 6/11 hyperparameter sets were capable of accurately learning all six tasks (>80% accuracy over the final 100 trials per task, averaged across the 5 seeds; Figure 4B), without backpropagation. RNNs from three of these hyperparameter sets achieved >85% accuracy averaged across all seeds, and RNNs from one hyperparameter set achieved >90% accuracy, demonstrating that this paradigm can support reliable and accurate learning of multiple tasks. RNNs from 3/11 hyperparameter sets achieved an intermediate level of performance (between 75 and 80% accuracy when averaged across seeds), successfully learning the tasks for some random seeds but not others. For the remaining 2/11 hyperparameter sets, performance was highly variable across random seeds – while some runs reached above 90%, on others the RNNs failed even to learn to fixate, yielding an accuracy of 0% (Figure 4C). All together, this experiment represents a proof-of-concept that interacting RL actors can learn cognitively demanding memory-based tasks without the need for backpropagation through hidden layers or through time.

## Discussion

In this study, we sought to determine whether rapid multitask learning can be accomplished despite limiting synaptic changes to highly localized sections of the network, the output and top-down layers, comprising ∼0.1% of all network connections. To do so, we generated thousands of different RNNs, each with a different combination of network properties that govern topology, normalization, etc., and found that only a handful of RNNs with specific network properties were capable of rapidly learning all tasks. Most RNNs generated using these high-performing network properties could reliably solve all tasks, and could generalize to novel tasks. To demonstrate that this form of learning could be performed in a more biologically-plausible manner, we modified our learning algorithm to use only local reward signals (i.e. without backpropagation), using a reinforcement learning setup involving two cooperating actors. This learning algorithm allows RNNs with high-performing network properties to learn all tasks. Our results suggest that biologically-inspired artificial networks, and potentially even biological organisms, are capable of rapidly learning multiple tasks with high localized synaptic plasticity using only local reward signals.

The idea that networks with (mostly) fixed connection weights can learn to perform different tasks was proposed over two decades ago, and is known as either echo state networks [24] or liquid state machines [25]. In both cases, a “reservoir” of randomly connected neurons with fixed connections weights respond to an input stimulus in a diverse and nonlinear manner. Neural responses within this network, in principle, form a basis set which can be linearly combined downstream to solve any task without requiring changes in synaptic weights within the network. A later variant known as FORCE training [26] demonstrated the importance of providing feedback onto the recurrent network. However, interest in these techniques has waned after backpropagation through time emerged as a powerful method for training RNNs [33]. Furthermore, training can only be accomplished using supervised learning, limiting the use of these methods to cases where a teaching signal is provided. However, our results suggest that the main idea of these studies–that random, fixed-connection RNNs are capable of solving complex tasks–is valid. The caveat is that it requires 1) that the RNNs are correctly initialized (which was already known) and 2) learning a task-dependent signal (a single linear transformation in this case) that projects onto the recurrent layer, allowing it to flexibly switch between tasks. Crucially, we show that this task-dependent signal can be learned in a biologically-plausible manner using only local error signals by structuring our networks as a system of two interacting actors trained using reinforcement learning.

This work also follows the spirit of the “lottery ticket hypothesis” [34], proposing that neural networks contain sparse, fortuitously-initialized subnetworks (so called ‘winning tickets’, akin to biological neural networks shaped by evolution and development). When trained in isolation, these subnetworks can reach performance equivalent to that obtained by the full model. Further studies demonstrated that this phenomenon generalizes to different network architectures and tasks [35–37]. These studies are consistent with our results suggesting that learning new tasks can be performed by networks with highly-limited plasticity.

Common to all of these studies is that learning new tasks with limited synaptic plasticity is dependent on proper network initialization. This tentatively suggests that one important role of neurodevelopment is the correct wiring of neural circuits so that they can learn future tasks with relatively sparse task-related synaptic plasticity. Specifically, one hypothesis is that synaptic plasticity, along with other molecular and cellular processes, initializes neural circuits to properly process and transmit information, allowing them to innately perform various computations without the need to alter synaptic connections [38]. Future work could build upon our growing understanding of circuit development in biological networks [39, 40] to determine whether RNNs equipped with the same developmental logic as their biological counterparts can learn to perform similar tasks.

In this study, the learned top-down signal that biases the recurrent layer plays a role similar to one hypothesized for subcortical circuits, such as the thalamus [41, 42], the basal ganglia [43], and the cerebellum [44]. For example, recent experimental studies have shown that the mediodorsal thalamus can encode the current behavioral context, allowing prefrontal cortex to flexibly switch representations during different contexts [41]. This might suggest that when an animal is confronted by a new task, context, or environment, synaptic plasticity might (initially) be located primarily in subcortical circuits to facilitate rapid learning [45], consistent with experimental evidence [9, 46]. Although the network architecture in our study only crudely approximates the role of subcortical structures implicated in the execution of context-dependent behaviors, we believe that by drawing inspiration from the neural circuits found *in vivo*, we can build artificial neural networks that better approximate the ability of humans and advanced animals to seamlessly switch between different contexts.

While this study demonstrates that networks with highly limited synaptic plasticity can rapidly learn new tasks in a biologically-plausible manner, it also leaves open questions related to initialization and robustness. We found that networks’ initial connectivity was the critical determinant of their ability to rapidly learn, and we relied on a random search to identify such hyperparameters. In reality, multiple processes shape the connectivity of neural circuits in the brain–some that are driven stochastically and act over long timescales, like evolution, and others that are more deterministic, like plasticity mechanisms that remodel connectivity during development. Modeling a broader spectrum of these biologically-anchored mechanisms could help encourage proper network connectivity in a more efficient and principled way. A similar approach of looking to biology for inspiration may also ameliorate issues of learning instability, as some of our high-performing and “reliable” RNNs failed to learn the tasks using RL. These networks might benefit from mechanisms that better regulate neural activity; improving our implementation of divisive normalization [47] (see Methods) or adding damping mechanisms such as spike rate adaptation [48] are two possible approaches. Finally, several recent studies have proposed additional mechanisms to facilitate biologically-plausible learning, such as adding dendrites to model neurons [18, 49], or using control theory to properly modulate the feedback onto a network [50]. Integrating these processes could further stabilize learning, and perhaps allow our networks to learn even more demanding tasks in a more biologically-plausible manner.

In summary, we have shown that networks can rapidly learn multiple tasks despite limiting synaptic changes to the output and top-down layers (comprising ∼ 0.1% of network connections). While this hypothesis must be experimentally verified, we hope that biologically-inspired machine learning models can continue to rapidly generate new hypotheses about neural circuits found *in vivo*, as well as test their plausibility. The close interplay between computational and experimental work in artificial networks as well as experimental model organisms provides a catalyst for novel biological insights, which in turn provide inspiration to create novel machine learning algorithms capable of more rapid, robust, and energy efficient learning.

## Methods

### Cognitive tasks

Networks were trained to perform two sets of cognitive tasks described below. The first six tasks (Figure 1A) all shared a common temporal structure, and were used to perform a large-scale sweep of network hyperparameters (Figure 2), and to test the RL-based learning algorithm (Figure 4). The second set of two tasks involved longer temporal dependencies, and was used to test how high-performing network hyperparameters generalize to novel tasks (Figure 3C). All tasks included the same core epochs: fixation, sample, delay, and test. The stimulus in all tasks was visual motion, with six possible directions, and was presented during the sample epoch (i.e. the sample stimulus), and possibly during the test epoch (i.e. the test stimulus).

For the delayed match-to-sample (DMS) task, the networks had to indicate whether the sample motion direction was an identical match to the test motion direction. For the anti-DMS task, networks had to indicate whether the test motion direction was 180° rotated from the sample motion direction.

For the delayed match-to-category (DMC) task, the networks had to indicate whether the sample motion direction was a categorical match to the test motion direction. Stimulus categories were defined by dividing the motion directions into two equal-sized groups — 0-180 degrees = category 1, 180-360 degrees = category 2. For the anti-DMC task, networks had to indicate whether the sample and test categories did not match (i.e. the correct response for the DMC and anti-DMC was reversed).

For the one interval categorization (OIC) task, the networks had to indicate whether the sample stimulus belonged to category 1 or category 2. For the anti-OIC task, the correct response was reversed. No test stimulus was presented.

For the two delayed match-to-sample tasks with multiple distractors (A-B-B-A and A-B-C-A), a sample stimulus was followed by three sequentially presented test stimuli (separated by delay epochs), and networks had to indicate whether each test motion direction matched the sample motion direction. In the A-B-B-A task, if a test motion direction was a non-match, there was a 50% probability that the test motion direction would be repeated immediately. In the A-B-C-A task, non-matching test stimuli were never repeated during a single trial.

For all tasks, the network were trained to maintain fixation during the fixation, sample and delay epochs, and then generate the correct response (as detailed above) during the test epoch(s). A trial was deemed correct if it generated the correct response during the test epoch, and maintained fixation for all prior time points. For the A-B-B-A and A-B-C-A tasks, accuracy was defined as the average accuracy across the three test epochs.

To train networks using reinforcement learning (Figure 4), we assigned rewards (both positive and negative) at each time step based on the network output. A correct decision during the test period earned a reward of 1.0, while an incorrect decision during the test period earned a reward of -0.01 (i.e. a penalty). Breaking fixation at any time point outside the test period earned a reward of -1.0, and failing to break fixation during the entire test period also earned a reward of -1.0. A reward of 0.0 was given in all other cases. A trial was terminated at any time if the network’s action resulted in a nonzero reward, whether positive or negative.

### Network model

RNNs were trained and simulated using the Python machine learning framework TensorFlow. The network architecture, including all layers and the connections between them, is schematized in Figure 1B when trained using supervised learning, and in Figure 4A when trained using reinforcement learning. All experiments used rate-based, biologically-inspired RNN models, similar to those previously described in [21–23]. Units in these networks obey Dale’s principle (e.g. either excite or inhibit all of their postsynaptic partners), and take on strictly positive, non-saturating activities obtained via the ReLU activation function.

The stimulus input layer consisted of 32 motion direction tuned neurons and 1 fixation tuned neuron. The tuning of the motion direction selective neurons followed a von Mises distribution, such that the activity of the input neuron *i* was

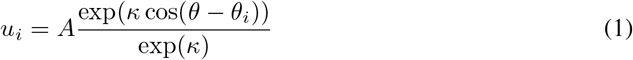

where *θ* was the direction of the motion stimulus, *θ*_*i*_ was the preferred direction of the neuron, *κ* was set to 2, and *A* was set to 2 when the motion direction stimulus was on (sample and test epochs) and set to 0 otherwise. The activity of the fixation neuron was set to 4 when the fixation cue was on (during the fixation, sample, and delay epochs), and was set to 0 when the fixation cue was off (test epoch).

The input weights were fixed (i.e. not trained) and projected onto half of the neurons in the recurrent layer (50% of the excitatory neurons and 50% of the inhibitory neurons). The top-down layer consisted of 6 neurons, encoding the 6 tasks using a one-hot representation. These 6 neurons projected onto a 64-dimensional vector via a trainable linear projection. This 64-dimensional vector was then projected onto half of the neurons in the recurrent network (the other half than those receiving input projections). The projection of the 64-dimensional top-down vector onto the recurrent layer was fixed.

The recurrent layer consisted of 2000 excitatory and 500 inhibitory neurons. To make it more challenging to maintain information in short-term memory, neither short-term synaptic plasticity [23] nor firing rate adaptation [16] was used.

### Network dynamics

The equations governing network dynamics are similar to those used in recent studies [21, 23]. We let **u** represent the activity of the stimulus input, **c** represent the activity of the context signal, and **r** represent the activity of the recurrent layer. The context signal is first linearly transformed into a top-down signal: **v** = **c***W*^*c*^, where the linear transformation *W*^*c*^ is a trainable matrix. The activity of the recurrent neurons was then modeled to follow the dynamical system [21, 23]:

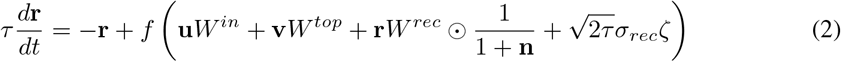

where *τ* is the neuron’s time constant, set to 100ms, *f* (·) is the ReLu activation function, *W* ^*in*^, *W* ^*top*^ and *W* ^*rec*^ are the input, top-down and recurrent connection weights, *ζ* is independent Gaussian white noise with zero mean and unit variance applied to all recurrent neurons and *σ*_*rec*_ is the strength of the noise. *σ*_*rec*_ was set to 0.05 when training using supervised learning, and to 0 when training with reinforcement learning. The connection weights *W* ^*in*^, *W* ^*top*^, and *W* ^*rec*^ are fixed (i.e. not trainable).

The vector **n** is a normalization term that controls the effective strength of recurrent connectivity [47]. It is governed by the system

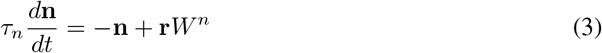

where *τ*_*n*_ is the normalization time constant, and the matrix *W* ^*n*^ determines how activity across the recurrent layers shunts the activity of each neuron.

The output layer consisted of a policy vector, which determines the action of the network, and when training using reinforcement learning, a critic, which predicts the discounted future reward. These are computed as

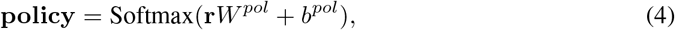

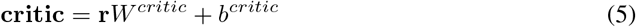

where *W* ^*pol*^, *W* ^*critic*^ are the connection weights and and *b*^*pol*^, *b*^*critic*^ are the biases. These four sets of parameters are trainable.

To simulate the RNN, we used a first-order Euler approximation with time step Δ*t*, which was set to 20ms:

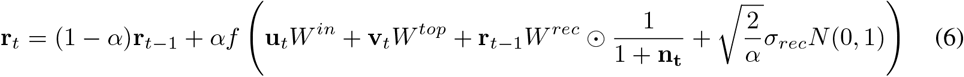

where 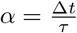 and *N* (0, 1) indicates the standard normal distribution. Normalization was similarly approximated as

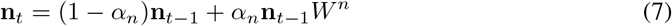

where 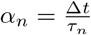

### Network properties and their random generation

Since we did not know *a priori* how to properly initialize RNNs capable of rapidly solving tasks with constrained plasticity, we developed a method to randomly generate RNNs from a set of 26 hyperparameters that were loosely modelled on the properties of neural circuits found *in vivo*. To generate a new RNN, we randomly sampled these 26 hyperparameters from uniform distributions with defined ranges (defined below). These 26 sampled values would then parameterize distributions from which we randomly sample the connections weights. We describe the 26 hyperparameters below.

We generated the initial recurrent weights in a series of steps. We first sampled the initial excitatory-to-excitatory, excitatory-to-inhibitory, inhibitory-to-excitatory, and inhibitory-to-inhibitory connections from separate gamma distributions, each with a scale parameter of 1.0 and a shape parameter of *κ*_*EE*_, *κ*_*EI*_, *κ*_*IE*_, *κ*_*II*_, respectively. Given that reciprocal connections occur more often than chance in cortex [51], we then increased the weights between reciprocally connected neurons, and decreased the weights of all non-reciprocally connected neurons as:

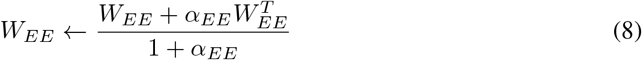

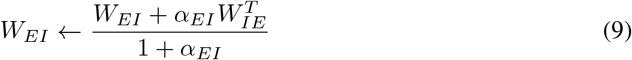

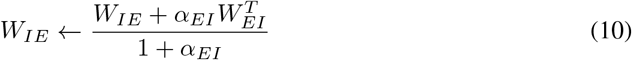

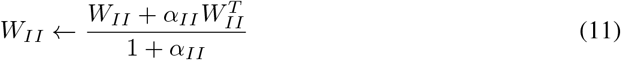

where ^*T*^ represents the transpose.

Given the presence of topographic maps in many cortical areas [52], we added topological structure by embedding the recurrent layer with a ring-like topology. Specifically, both excitatory and inhibitory neurons were assigned angles equally distributed between 0 and 2*π*. We then increased connection weights between neurons that were close together according to the cosine of their angular distance, cos *θ*, and decreased connection weights between those neurons that were far apart using a von Mises distribution:

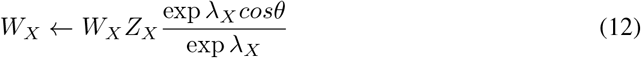

where X represents either *EE, EI, IE*, or *II*, and *Z*s are chosen such that the mean connection weight for each group is left unchanged. Finally, we scaled all connection weights by a global scalar *w*_*r*_.

Bottom-up and top-down weights were initialized in a similar manner. The initial connection weights were sampled from gamma distributions where *κ*_*BU,E*_, *κ*_*T D,E*_ were the shape hyperparameters of bottom-up and top-down projections onto the recurrent excitatory neurons, and *κ*_*BU,I*_, *κ*_*TD,I*_ the shape hyperparameters of bottom-up and top-down projections onto the recurrent inhibitory neurons. Both bottom-up and top-down neurons were also endowed with a topological structure by evenly spacing them along a ring, in which they were assigned angles equally distributed between 0 and 2*π*. For bottom-up neurons, this angle was identical to their preferred motion direction. We then applied the same Von Mises-like function as in Equation 12 to scale weights based upon their angular distance. This was done separately for the bottom-up and top-down projections onto excitatory neurons using λ_*BU,E*_ and λ_*TD,E*_, respectively, and for the bottom-up and top-down projections onto inhibitory neurons using λ_*BU,I*_ and λ_*T D,I*_, respectively. Finally, we also scaled all bottom-up weights by a global scalar *w*_*BU*_.

**Table 1:** Network weight initialization hyperparameters

| Recurrent weight distribution shape/topology |  |  |  |
| --- | --- | --- | --- |
|  | *Name* | *Description* | *Range* |
| 1 | $\kappa_{EE}$ | Shape of excitatory-to-excitatory (EE) weight distrib. | [0.05, 0.25] |
| 2 | $\kappa_{EI}$ | Shape of excitatory-to-inhibitory (EI) weight distrib. | [0.05, 0.25] |
| 3 | $\kappa_{IE}$ | Shape of inhibitory-to-excitatory (IE) weight distrib. | [0.05, 0.25] |
| 4 | $\kappa_{II}$ | Shape of inhibitory-to-inhibitory (II) weight distrib. | [0.05, 0.25] |
| 5 | $\lambda_{EE}$ | Shape of topological modifier for EE connectivity | [0, 2] |
| 6 | $\lambda_{EI}$ | Shape of topological modifier for EI connectivity | [0, 2] |
| 7 | $\lambda_{IE}$ | Shape of topological modifier for IE connectivity | [0, 2] |
| 8 | $\lambda_{II}$ | Shape of topological modifier for II connectivity | [0, 2] |
| 9 | $w_r$ | Global strength of recurrent weights | [0.05, 0.2] |
| Recurrent reciprocal connectivity |  |  |  |
| 10 | $\alpha_{EE}$ | Strength of reciprocal connectivity between pairs of E units | [0, 0.3] |
| 11 | $\alpha_{EI}$ | Strength of EI/IE reciprocal connectivity | [0, 0.3] |
| 12 | $\alpha_{II}$ | Strength of II reciprocal connectivity | [0, 0.3] |
| Bottom-up input weight distribution shape/topology |  |  |  |
| 13 | $\kappa_{BU,E}$ | Shape of bottom-up input weight distribution onto E units | [0.05, 0.25] |
| 14 | $\kappa_{BU,I}$ | Shape of bottom-up input weight distribution onto I units | [0.05, 0.25] |
| 15 | $\lambda_{BU,E}$ | Shape of topological modifier for bottom-up input onto E units | [0, 2] |
| 16 | $\lambda_{BU,I}$ | Shape of topological modifier for bottom-up input onto I units | [0, 2] |
| 17 | $w_{BU}$ | Global strength of bottom-up input weights | [0.05, 0.2] |
| Activity normalization weights |  |  |  |
| 18 | $\beta_{EE}$ | Strength of normalization between pairs of E units | [0, 2] |
| 19 | $\beta_{EI}$ | Strength of normalization from E to I units | [0, 2] |
| 20 | $\beta_{IE}$ | Strength of normalization from I to E units | [0, 2] |
| 21 | $\beta_{II}$ | Strength of normalization from I to I units | [0, 2] |
| 22 | $\tau_n$ | Time constant of network activity modulation | [20, 100] |
| Top-down input weight distribution shape/topology |  |  |  |
| 23 | $\kappa_{TD,E}$ | Shape of top-down weight distribution onto E units | [0.05, 0.25] |
| 24 | $\lambda_{TD,E}$ | Shape of topological modifier for top-down input onto E units | [0.05, 0.25] |
| 25 | $\kappa_{TD,I}$ | Shape of top-down weight distribution onto I units | [0, 2] |
| 26 | $\lambda_{TD,I}$ | Shape of topological modifier for top-down input onto I units | [0, 2] |

The matrix that determines how activity in the recurrent layer normalizes activity, *W* ^*n*^, consisted of four unique values:

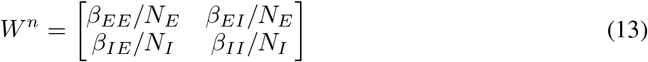

where *N*_*E*_ and *N*_*I*_ were the number of excitatory (2000) and inhibitory (500) neurons in the recurrent layer, respectively. The time constant that controls the dynamics of normalization in Equation 3 was defined as *τ*_*n*_.

Finally, the bias term for all neurons was initialized to 0. All self-connections were set to 0, and all initial weights above 2 were clipped.

### Network architecture for reinforcement learning of cognitive tasks

When training using reinforcement learning, our network was comprised of two different actors. The “external” actor, which consisted of the output layer, was responsible for selecting discrete actions to solve the cognitive tasks. Its policy vector was of size 3, and the critic vector was of size 2 (described below). The “internal” actor, which consisted of the top-down layer, was responsible for selecting continuous action signals that were projected onto the recurrent layer. Its policy vector was of size 64, and actions were sampled from a normal distribution with a mean value equal to the policy vector, and a standard deviation that was initially set to 0.1 at the start of training and then linearly decayed to 0.01 by the end of training. The sampled actions were capped at *±* 5.

The 2-D critic vector was responsible for predicting the discounted future reward at two different timescales. The fast timescale, with a discount factor *γ* = 0.9, was used to train the external (discrete) actor, and the slow timescale, with a discount factor *γ* = 0.95, was used to train the internal (continuous) actor.

### Network training and testing

For supervised learning, we used the cross-entropy loss function. Network weights were trained using the Adam optimizer [53] with the learning rate set to 0.02 for the output layer, 0.002 for the top-down layer, and *ϵ* = 10^−7^. The batch size was 256.

When training the two cooperating actors using reinforcement learning, we used the proximal policy optimization (PPO) algorithm [54]. Network weights were also trained using the Adam optimizer, with a learning rate of 5 × 10^−4^ for the external actor (i.e. the policy and critic layers), 5 × 10^−6^ for the internal actor (i.e. the top-down layer), and epsilon was set to 10^−5^. The batch size was 256, the time horizon was 16 time steps, and training data was split into 4 mini-batches. The external actor was trained for 3 epochs, while the internal actor was trained for 1 epoch. The clip ratio in both cases was 0.1. Finally, we used the generalized advantage estimate [55] for the internal actor, with λ = 0.9. The advantage estimate for both the external and internal actors was not normalized.

### Measuring the strength of motion encoding during the delay period

We quantified how well the RNNs could maintain sample information in short-term memory by measuring how accurately we could decode the motion direction using multiclass support vector machines (SVMs). Since we were interested in how well RNNs could innately maintain information in short-term memory, this calculation was performed before the RNNs were trained on any tasks. We used multiclass SVMs to classify the motion direction using the neuronal activity from the 2500 recurrent neurons measured at the end of the delay period from a batch of 512 trials. We measured the classification accuracy using 4-fold stratified cross-validation, in which we randomly split the data into 4 folds, while ensuring that all folds contained roughly even proportions of trials with each of the 6 possible stimuli. We then fit an SVM with a linear kernel on each trio of folds (75% of trials) and tested its performance on trials in the fourth held-out fold (25% of trials). We report the average test classification performance across all 4 train/test splits.

## Supplementary Information

To determine whether training the top-down weights was necessary for networks to accurately learn all tasks, we repeated our sweep used in Figure 2, except that top-down weights were randomized across tasks but frozen from initialization, and only output weights were trained. We trained 2,725 RNNs with fixed top-down weights (same as in original sweep), and mean accuracy after training was significantly less compared to RNNs in which the top-down weights were trainable (*p <* 10^−24^, Wilcoxon rank-sum test, Figure S1B).

Although the mean accuracy of RNNs with fixed top-down weights was reduced, it was still possible that a few RNNs were capable of accurately learning all tasks. Thus, we compared the top 20 hyperparameter sets from both sweeps, and resampled and retrained RNNs from each hyperparameter set 5 times (Figure S1A). Consistent with above, both the mean accuracy across the six tasks (*p <* 10^−7^, Wilcoxon rank-sum test) and the minimum accuracy (*p <* 10^−7^) was lower for RNNs with fixed top-down weights, confirming that networks with trainable top-down weights are more capable of rapidly learning multiple tasks.

**Figure S1:**
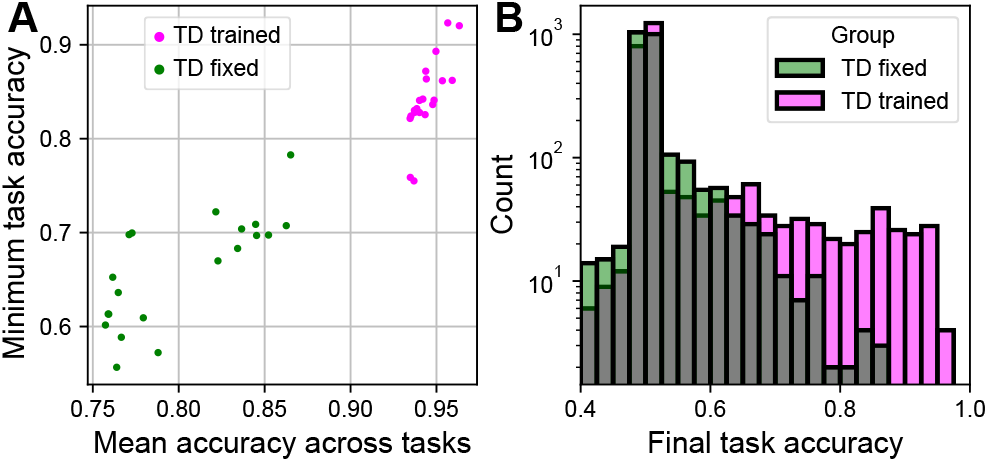
Fixing top-down inputs degrades learning. **A)** Scatter plot showing mean vs. minimum accuracy across the six tasks, averaged across 5 seeds of network weights. Magenta circles are from the 20 top-performing hyperparameter sets with top-down weights were trainable; green circles are from the 20 top-performing hyperparameter sets, obtained in a separate sweep for networks with top-down weights were fixed. **B)** Histogram of average accuracy across all six tasks at the end of training for 2 separate sweeps through hyperparameter space (magenta: top-down weights were trainable, green: top-down weights were fixed). Overlapping regions between the two sweeps are shown in grey. y-axis is shown in log-scale.

